# HebbPlot: An intelligent tool for learning and visualizing chromatin mark signatures

**DOI:** 10.1101/207670

**Authors:** Hani Z Girgis, Alfredo Velasco, Zachary E Reyes

## Abstract

Histone modifications play important roles in gene regulation, heredity, imprinting, and many human diseases. The histone code is complex, consisting of about 100 marks. Biologists need computational tools for characterizing general signatures representing the distributions of tens of chromatin marks around thousands of regions. To this end, we developed a software tool called HebbPlot, which utilizes a Hebbian neural network to learn such signatures. HebbPlot presents a signature as a digitized image, which can be easily interpreted. We validated HebbPlot in six case studies. HebbPlot is applicable to a wide array of studies, facilitating the deciphering of the histone code.

## Background

Understanding the effects of histone modifications will provide answers to important questions in biology and will help with finding cures to several diseases including cancer. Carey highlights several functions of epigenetic factors including Cytosine methylation and histone modifications [1]. It was reported that methylation of CpG islands inhibit transcription [2], whereas the complex histone code has a wide range of regulatory functions [3,4]. Additionally, epigenetic marks may affect body weight and metabolism [5]. Interestingly, chromatin marks may explain how some acquired traits, such as obesity and due to exposure to some toxins, are passed from one generation to the next (Lamarckian inheritance) [6–9]. Further, epigenetics may explain how two identical twins have different disease susceptibilities [10]. Epigenetic factors play a role in imprinting, in which a chromosome, or a part of it, carries a maternal or a paternal mark(s) [11,12]. Defects in the imprinting process may lead to several disorders [13–18], and may increase the “birth defects” rate of assisted reproduction [19]. Furthermore, chromatin marks play a role in cell differentiation by selectively activating and deactivating certain genes [20,21]. Some chromatin marks take part in deactivating one of the X chromosomes [22]. It has been observed in multiple types of cancer that some tumor suppressor genes were deactivated by hyperme-thylating their promoters [23–25], the removal of activating chromatin marks [26,27], or adding repressive chromatin marks [28]. Utilizing such knowledge, anticancer drugs that target the epigenome [1] have been designed. Two compounds are used in these drugs. One compound inhibits DNA methylation [29,30], whereas the other compound inhibits histone deacetylation [31] (histone acetylation is an activating mark).

Pioneering computational and statistical methods for deciphering the histone code have been developed. Some tools are designed for profiling and visualizing the distribution of a chromatin mark(s) around multiple regions [32,33]. Additionally, a tool for clustering and visualizing genomic regions based on their chromatin marks has been developed [34]. Several systems are available for characterizing histone codes/states in an epigenome [35–43]. Further, an alphabet system for histone codes was proposed [44]. Other tools can recognize and classify the chromatin signature associated with a specific genetic element [45–55]. Furthermore, methods that compare the chromatin signature of healthy and sick individuals are currently available [56].

Scientists have identified about 100 histone marks [37]. Additionally, there will be a near infinite number of future studies, in which scientists need to characterize the pattern of chromatin marks around a set of regions in the genome. Therefore, there is a definite need for an automated framework that enables scientists to (i) automatically characterize the chromatin signature of a set of sequences that have a common function, e.g. coding regions, promoters, or enhancers; and (ii) visualize the identified signature in a simple intuitive form. To meet this need, we designed and developed a software tool called HebbPlot. This tool allows average users, without extensive computational knowledge, to characterize and visualize the chromatin signature associated with a genetic element automatically.

HebbPlot includes the following four innovative approaches in an area that has become the frontier of medicine and biology:

- HebbPlot can learn the chromatin signature of a set of regions automatically. Sequences that have the same function in a specific cell type are expected to have similar marks. The learned signature represents these marks around all of the regions. *HebbPlot differs from the other tools in its ability to learn one signature representing the distributions of all available chromatin marks around thousands of regions*.
- This is the first application of Hebbian neural networks in the epigenetics field. These networks are capable of learning associations; therefore, they are well suited for learning the associations among tens of marks and genetic elements.
- The framework enables average users to train artificial neural networks *automatically.* Users are not burdened with the training process. Self-trained systems for analyzing protein structures and sequence data have been proposed [57–59]. HebbPlot is the analogous system for analyzing chromatin marks.
- HebbPlot is the first system that integrates the tasks of learning and visualizing a chromatin signature. Once the signature is learned, the marks are clustered and displayed as a digitized image. This image shows one pattern representing thousands of regions. To illustrate, the distributions of the marks appear around one region; however, they are learned from all input regions.

We have applied our tool to learning and visualizing the chromatin signatures of several active and inactive genetic elements in 57 tissues/cell types. These case studies demonstrate the applicability of HebbPlot to many interesting problems in molecular biology, facilitating the deciphering of the histone code.

## Materials and methods

In this section, we describe the computational principles of our software tool, HebbPlot. The core of the tool is an unsupervised neural network, which relies on Hebbian learning rules.

### Region representation

To represent a group of histone marks overlapping a region, these marks are arranged according to their genomic locations on top of each other and the region. Then equally-spaced vertical lines are superimposed on the stack of the marks and the region. The numerical representation of this group of marks is a matrix. A row of the matrix represents a mark. A column of the matrix represents a vertical line. If the *i^th^* mark intersects the *j^th^* vertical line, the entry i and j in the matrix is 1, otherwise it is −1. The first vertical line is at the beginning of the region; the last vertical line is at the end of the region; the rest of the lines are spread out evenly. Fig 1 shows the graphical and the numerical representations of a region and the overlapping marks. Finally, the two-dimensional matrix is converted to a one dimensional vector called the epigenetic vector. The number of vertical lines is determined experimentally. We used 41 and 91 lines in our experiments. This number should be adjusted according to the average size of a region.

**Figure 1.**
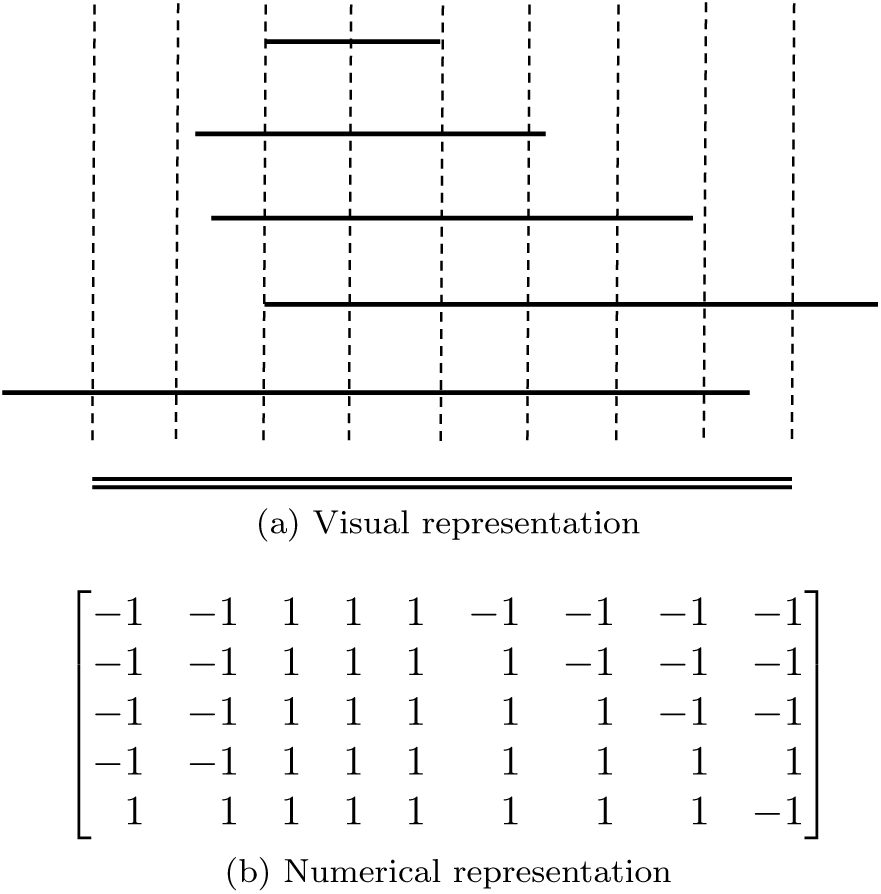
Representations of a group of chromatin marks overlapping a region. (a) Horizontal double lines represent a region of interest. Horizontal single lines represent the marks. Vertical lines are spaced equally and bounded by the region. (b) The intersections between the marks and the vertical lines are encoded as a matrix where rows represent the marks and columns represent the vertical lines. If a vertical line intersects a mark, the corresponding entry in the matrix is 1, otherwise it is −1.

### Data preprocessing

Preprocessing input data is a standard procedure in machine learning. During this procedure, the noise in the input data is reduced. First, vectors that consist mainly of −1’s are removed. These regions are very likely false positives. Then, each epigenetics vector is compared to two other vectors selected randomly from the same set. The value of an entry in the vector is kept if it is the same in the three vectors, otherwise it is set to zero. For example, consider the vector [1 1 −1]. Suppose that the vectors [1 −1 −1] and [1 −1 −1] were selected randomly. The result would be [1 0 −1] because the first and the third elements are the same in the three vectors, but the second element is not.

### Hebb’s network

Associative learning, also known as Hebbian learning, is inspired by biology. “When an axon of cell A is near enough to excite a cell B and repeatedly or persistently takes part in firing it, some growth process or metabolic change takes place in one or both cells such that A’s efficiency, as one of the cells firing B, is increased” [60]. In behavioral psychology, Ivan Pavlov conducted a famous experiment, which demonstrated learning by association. In this experiment, a dog was trained to associate the sound of a bell with food; this dog salivated when it heard the bell whether or not food was present. The presence of food is referred to as the unconditioned stimulus, *p*^0^, and the sound of the bell is referred to as the conditioned stimulus, *p*. Associating these two stimuli together is the goal. After training, the response to either the conditioned stimulus or the unconditioned one is the same as the response to both stimuli combined [61].

In the context of epigenetics, a Hebbian network can be viewed as the dog in Pavlov’s experiment. The unconditioned stimulus, *p*^0^, is a one-dimensional vector representing the distributions of histone marks over a sequence e.g. one tissue-specific enhancer. This vector is referred to as the epigenetic vector; it is obtained as outlined earlier in this section. The conditioned stimulus is always the one vector, which include ones in all entries. We would like to train the network to give a response, analogous to the salivation of the dog, when it is given the ones vector, whether or not the epigenetic vector is provided. The response of the network is a prototype/signature representing the distributions of histone marks over the entire set of genomic locations, e.g. all enhancers of a specific tissue.

Eq 1 and Eq 2 define how the response of a Hebbian network is calculated. The training of the network is given by Eq 3 [61].

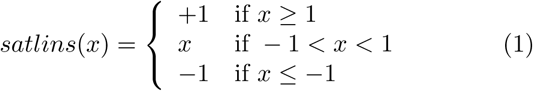

Eq 1 defines a transformation function. This function ensures that the response of the network is similar to the unconditioned stimulus, i.e. each element of the response is between 1 and −1. If x is a vector, the function is applied component wise.

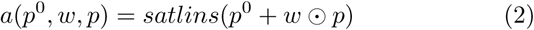

Eq 2 describes how a Hebbian network responds to the two stimuli. The response of the network is transformed using Eq 1. In Eq 2, *p*^0^ is the unconditioned stimulus, e.g. presence of food or an epigenetic vector; *W* is the weights vector, which is the prototype/signature learned so far; and *p* is the conditioned stimulus, e.g. sound of a bell or the one vector. The operator ⊙ represents the component-wise multiplication of two vectors. In the current adaptation, if the network is presented with an epigenetic vector and the one vector, the response is the sum of the prototype learned so far and the epigenetic vector. In the absence of the epigenetic vector, i.e. all-zeros *p*^0^, the response of the network is the prototype, demonstrating the ability of the network to learn associations.

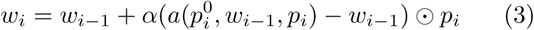

Eq 3 defines Hebb’s unsupervised learning rule. Here, *w_i_* and *w_i–1_* are the prototype vectors learned in iterations *i* and *i* – 1. The *i^th^* pair of unconditioned and conditioned stimuli is 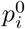 and *p_i_*. Learning occurs, i.e. the prototype changes, only when the *i^th^* conditioned stimulus, *p_i_*, has non-zero components. This is the case here because *p_i_* is always the one vector. Due to a small α, which represents the learning and the decay rates, the prototype vector changes a little bit in each iteration when learning occurs; it moves closer to the response of the network to the *i^th^* pair of stimuli.

### Comparing two signatures

One of the main advantages of the proposed method is that two signatures can be compared quantitatively. The dot product of two vectors indicates how close they are to each other in space. When these vectors are normalized, i.e. each element is divided by the vector norm, the dot product is between 1 and −1. The dotsim function (Eq 4) normalizes the vectors and calculates their dot product.

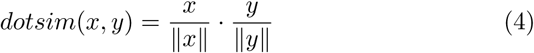

Here, x and y are vectors; ||*x*|| and ||*y*|| are the norms of these vectors; the · symbol is the dot product operator. It is easy to interpret the meaning of the dot product of two normalized vectors. If the two vectors are very similar to each other, the value of the dotsim function approaches 1. If the values at the same index of the two vectors are opposite of each other, i.e. 1 and −1, the value of dotsim approaches −1. The dotsim function can be applied to the whole epigenetic vector or to the part representing a specific chromatin mark. When comparing the chromatin signatures of two sets of regions, a mark with a dotsim value approaching 1 is common in the two signatures. A mark with a dotsim value approaching −1 has opposite distributions, distinguishing the signatures. Marks with dotsim values approaching zero do not have consistent distribution(s) in one or both sets; these marks should not be considered while comparing the two signatures.

### Visualizing a chromatin signature

Row vectors representing different marks are clustered according to their similarity to each other. We used hierarchical clustering in grouping marks with similar distributions. Hierarchical clustering is an iterative bottom-up approach, in which the closest two items/groups are merged at each iteration. The algorithm requires a pair-wise distance function and a cluster-wise distance function. For the pair-wise distance function, we utilized the city block function to determine the distance between two vectors representing marks. For the group-wise distance function, we applied the weighted pair group method with arithmetic mean [62]. A digitized image represents the chromatin signature of a genetic element. A one-unit-by-one-unit square in the image represents an entry in the matrix representing the signature. A row of these squares represents one mark. The color of a square is a shade of gray if the entry value is less than 1 and greater than −1; the closer the value to 1 (-1), the closer its color to white (black).

Up to this point, we discussed the computational principles of our software tool, HebbPlot. Next, we illustrate the data used in validating the tool.

### Data

We used HebbPlot in visualizing chromatin signatures characterizing multiple genetic elements. Specifically, we applied HebbPlot to:

1. Active promoters;
2. Active promoters on the positive strand;
3. Active promoters on the negative strand;
4. High-CpG active promoters;
5. Low-CpG active promoters;
6. Active enhancers;
7. Coding regions of active genes;
8. Coding regions of inactive genes; and
9. Random genomic locations.

The Roadmap Epigenomics Project provides tens of marks for more than 100 tissues/cell types [63]. Active genes were determined according to gene expression levels, which were obtained from the Expression Atlas [64] and the Roadmap Epigenomics Project [65]. A gene of expression level greater than 1 is considered active, whereas inactive genes are those having expression levels of 0. The coding regions were obtained from the University of California Santa Cruz Genome Browser [66]. The Ensemble genes for the hg19 human genome assembly were used in this study. Active promoters are those associated with active genes. A promoter region is defined as the 400-nucleotides-long region centered on the transcription start site. To divide the promoters into high- and low-CpG groups, we calculated the CpG content according to the method described by Saxonov, et al. [67]. Enhancers active in H1 and IMR90 were obtained from a study by Rajagopal, et al. [54]; this study provides the P300 peaks. We considered the enhancers to be the 400-nucleotides-long regions centered on the P300 peaks. Regions of enhancers active in liver, foetal brain, foetal small intestine, left ventricle, lung, and pancreas were obtained from the Fantom Project [68].

Once the locations of a genetic element were determined, they are processed further. If the number of the regions, e.g. tissue-specific enhancers, was more than 10,000 regions, we sampled uniformly 500 regions from each chromosome. Each region was expanded by 10% on each end to study how chromatin marks differ from/resemble the surrounding regions; except promoters on the positive and the negative strands. They were expanded by 500% on each end. Overlapping regions, if any, were merged.

### Availability

The software and the data sets are available as Additional files 1–8. Additionally, HebbPlot can be downloaded from GitHub (https://github.com/TulsaBioinformaticsToolsmith/HebbPlot).

In this section, we discussed the computational method and the data. Next, we apply HebbPlot in six case studies.

## Results

### Case study: Signature of H1-specific enhancers

We studied multiple enhancers active in the H1 cell line (human embryonic stem cells) obtained from a study conducted by Rajagopal, et al. [54]. These enhancers were detected using P300 ChIP-Seq. An enhancer is represented as a 400-nucleotides-long region centered on the P300 peak. This data set contains 5899 enhancers and 27 histone marks. Each enhancer region was expanded by 10% on each end to study how chromatin marks differ from/resemble the surrounding regions. To begin, 41 uniform samples/points were obtained from each region. Then for each point, it was determined whether or not it falls in a mark region overlapping the enhancer. Next, we plotted tens of these enhancers; four of these plots are shown in Figures 2a-2d. No clear signature appears in these plots. After that, a HebbPlot representing the signature of H1-specific enhancers was generated (Figure 2e) using an unsupervised hebbian network. For comparison purposes, we generated a conventional plot (Figure 2f).

**Figure 2.**
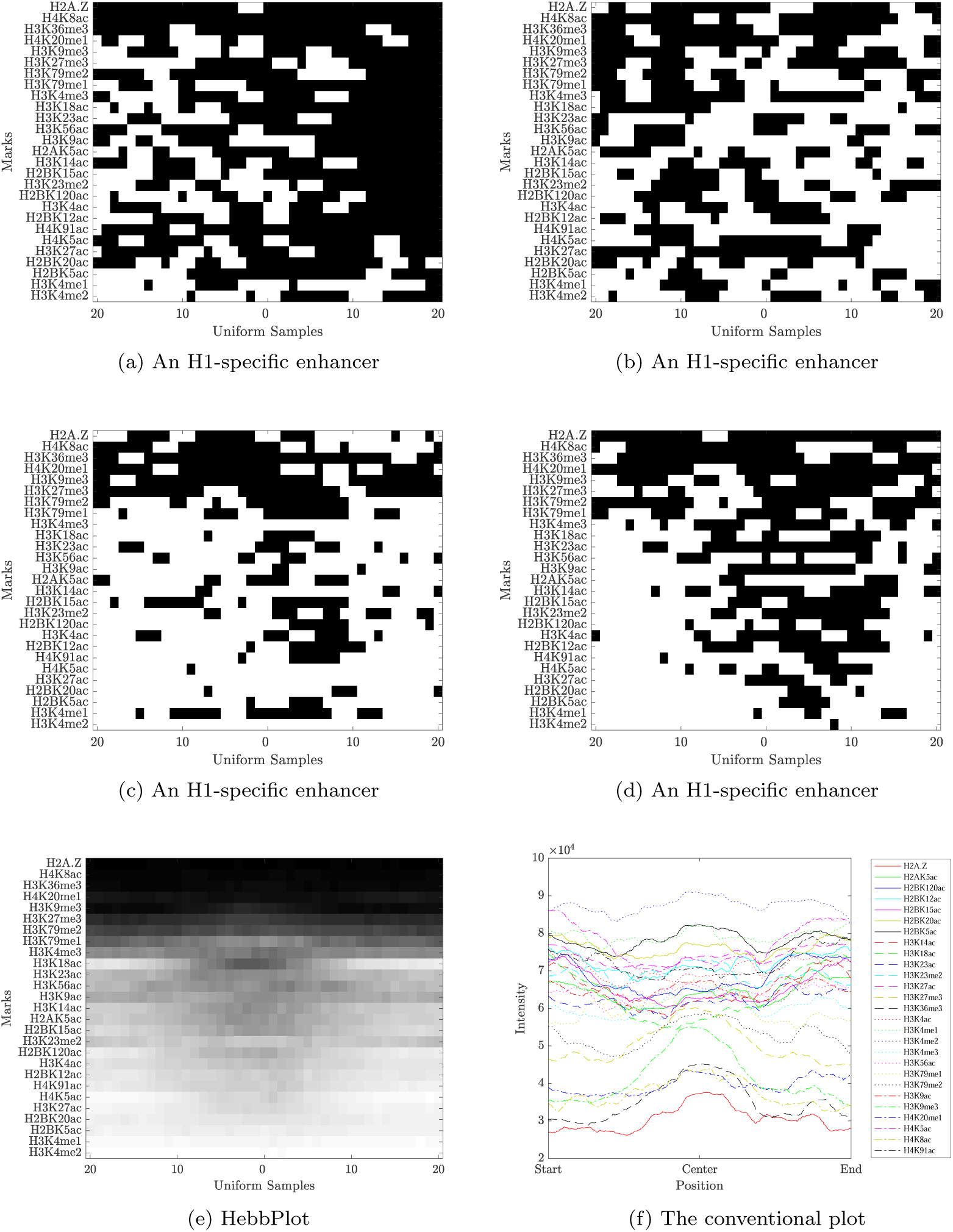
Retrieving the chromatin signature of the H1-specific enhancers. Four examples of enhancers are shown in Parts a–d. A row in one of these plots represents the distribution of one mark around a region; white (black) color indicates the presence (absence) of a mark. It is hard to see a common pattern in these four examples. The signature learned by the Hebbian network is captured by the HebbPlot shown in Part e. *A row in the HebbPlot represents the distribution of a mark around all enhancers in the data set*. The brighter the color, the higher the certainty of the presence of a mark around the corresponding sub-region. The HebbPlot is characterized by four zones. The top most zone represents chromatin marks that are absent from the enhancer regions, whereas the next three zones represent the present marks with increasing certainty. A conventional plot of the intensities of all marks around every region in the data set in shown in Part f. Many marks show depressions near the center of the plot; however, some peaks are mixed with these depressions in the conventional plot. In contrast, these depressions correspond to the dark ellipse in the middle of the third zone of the HebbPlot. This ellipse is very clear. Further, marks of similar intensities obstruct one another in the conventional plot. This is not the case with HebbPlot because every mark is represented by a separate row.

The HebbPlot shows four zones representing the absent marks, and the present ones with different confidence levels. For example, the top zone shows four marks (H2A.Z, H4K8ac, H3K36me3, and H4K20me1) that are absent from the H1 enhancers. The second zone from the top shows marks with very weak intensities including H3K9me3, H3K27me3, H3K79me2, and H3K79me1. The third zone has an ellipse, which is darker than the surrounding area, indicating that the signals of the marks within the ellipse are weaker than the surroundings. The bottom zone shows two marks (H3K4me1 and H3K4me2) that are present around these enhancers with the highest confidence level.

In the upper part of the conventional plot, a large number of marks show depressions near the middle of the plot. However, these depressions are mixed with few peaks, making viewing them hard. These depressions correspond to the fragments near the centers of the individual plots and the dark ellipse in the middle of the third zone of the HebbPlot. Clearly, the dark ellipse captures this pattern much better than the conventional plot. Further, marks with similar intensities overlap each other in the conventional plot, obstructing on another. The more the marks, the worse the obstruction. To illustrate, this figure was generated using 27 marks; there are about 100 known histone marks; therefore, using these conventional figures may not be the best way to visualize the intensities of a large number of marks. In contrast, HebbPlot can handle a large number of marks efficiently because each mark has its own row. Furthermore, no noise-removal process was applied while constructing the conventional figure. In contrast, only regions, or sub-regions, that are recognized by the network contribute to the HebbPlot.

### Case study: Histone signatures of different active elements in liver

Seven histone marks of the human liver epigenome are available. We obtained 5005 enhancers, 13688 promoters, and 12484 coding regions of active genes in liver. In addition, we selected 10,000 locations sampled uniformly from all chromosomes of the human genome as controls. Then we trained four Hebbian networks to learn the chromatin signature of each genetic element. As expected, the HebbPlot representing the random genomic locations displays a black box (not shown), indicating that no chromatin mark is distributed consistently around these regions. Figure 3 shows three HebbPlots of the enhancers, the promoters, and the coding regions. The three signa tures have similarities and differences. Two marks, H3K9me3 and H3K27me3, are absent from the three signatures. However, the three signatures are distinguishable. H3K36me3 is the strongest mark of the coding regions, whereas it is absent from the promoters and the enhancers. On the other hand, H3K27ac is the strongest mark on the promoters and the enhancers, but almost absent from the coding regions. H3K4me1 is stronger than H3K4me3 around the enhancers, but H3K4me3 is stronger than H3K4me1 around the promoters. Both of these marks are absent from the coding regions. These plots demonstrate that HebbPlot is able to learn the chromatin signature from a group of regions with the same function. In addition, the chromatin signatures of the promoters, the enhancers, and the coding regions have similarities and differences.

**Figure 3.**
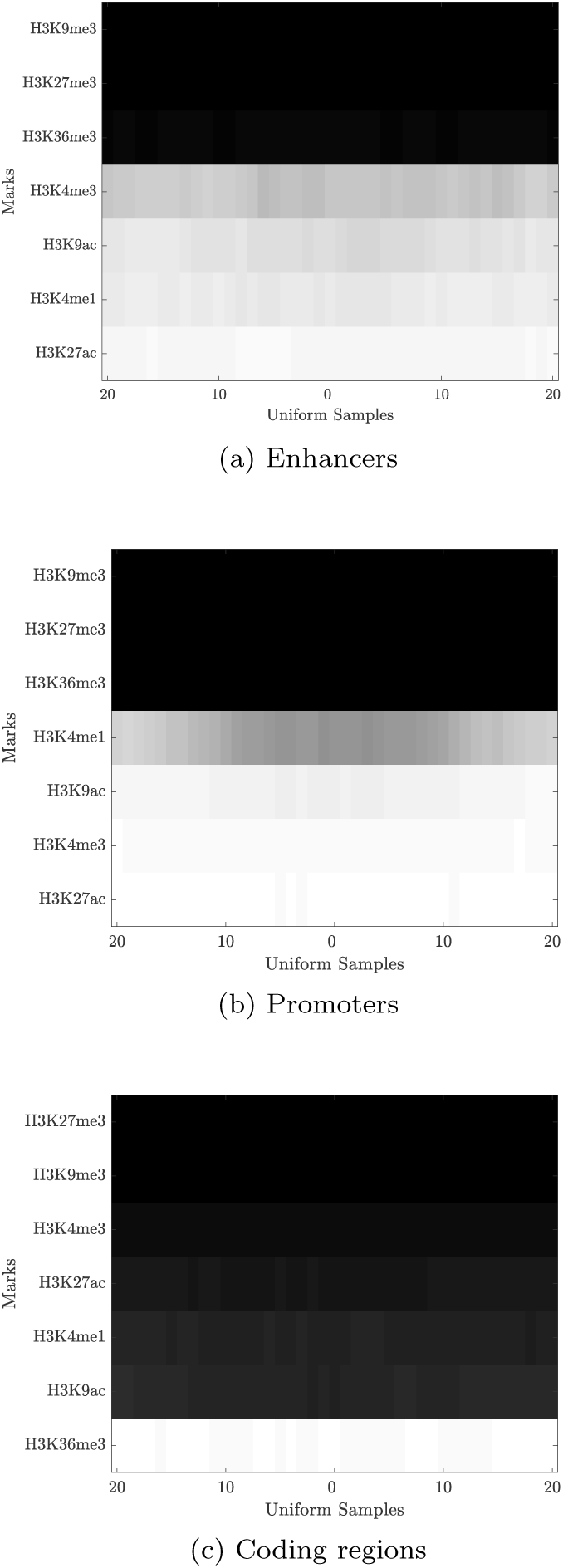
Liver chromatin signatures representing (a) active enhancers, (b) active promoters, and (c) coding regions of active genes. The three signatures have similarities and differences. They are similar in that H3K9me3 and H3K27me3 are absent from all of them. H3K36me3 is the strongest mark of coding regions, whereas H3K27ac is the strongest mark of promoters and enhancers. H3K4me1 is stronger than H3K4me3 in enhancers; this relation is reversed in promoters, where H3K4me1 is weak around transcription start sites.

### Case study: The directional signature of active promoters

Because promoters are upstream from their genes, some marks may indicate the direction of the transcription. To determine whether or not marks have direction, active promoters were separated according to the positive and the negative strands into two groups. Then the promoter region was expanded five times on each end. The expanded region has these three parts: (i) the 2000-nucleotides-long region upstream from a promoter, (ii) the 400-nucleotides-long promoter region itself, and (iii) the 2000-nucleotides-long region downstream from the promoter. We trained two Hebbian networks to learn the chromatin signatures of active promoters on the positive and the negative strands. Figure 4 shows the HebbPlots of the positive and the negative promoters active in HeLa-S3 cervical carcinoma cell line. These two plots are mirror images of each other, showing H3K36me3, H3K79me2, H3K4(me1,me2,me3), H3K27ac, and H3K9ac stretching more downstream than upstream and H2A.Z in the opposite direction.

**Figure 4.**
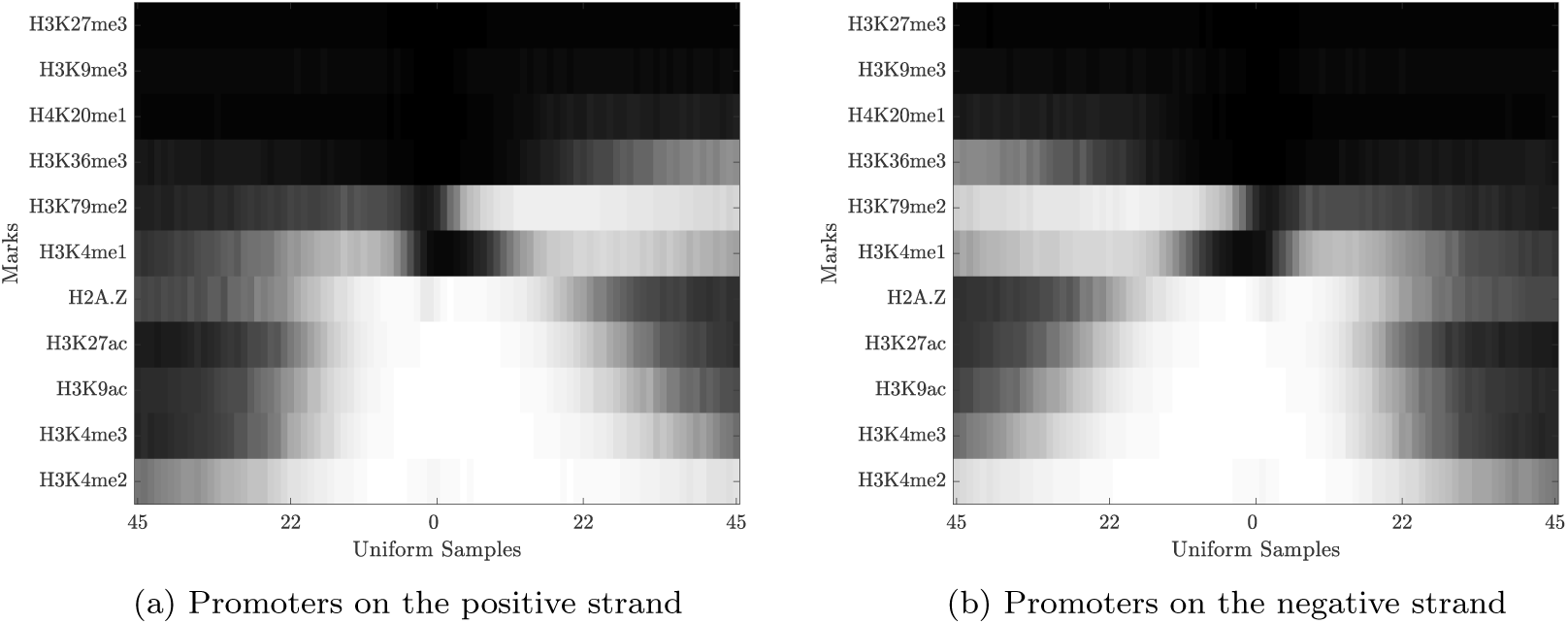
HebbPlots of active promoters in HeLa-S3 cervical carcinoma cell line. These promoters were separated into two groups according to their strands. Each promoter region was expanded five times on each end to investigate the directions of histone marks around these regions. The size of the expanded region is 4400 nucloetides. The two HebbPlots of the promoters on the positive and the negative strands are mirror images of each other. Multiple marks including H3K36me3, H3K79me2, H3K4me1, H2A.Z, H3K27ac, H3K9ac, H3K4me3, and H3K4me2 are distributed in a direction specific way. H2A.Z tends to stretch upstream, whereas the rest of these directional marks tend to stretch downstream from the promoters toward their coding regions.

Then we generated HebbPlots for the positive (Additional file 2) and the negative (Additional file 3) promoters of 57 tissues, for which we know their gene expression levels. The directional signature of promoters is very consistent in these tissues. After that, we determined quantitatively which marks having directional preferences in the 57 tissues/cell types. Recall that two vectors pointing in opposite directions have a dotsim value of −1. The closer the value to −1 is, the closer the angle between the two vectors to 180° is. To determine directional marks, the learned prototype of a mark over the upstream third of the expanded promoter region was compared to the prototype of the same mark over the downstream third. If the dotsim value between the two prototypes is negative, this mark is considered directional. We list the number of times a chromatin mark was determined for a tissue and the number of times it showed directional preference in Table 1. H3K4me3 and H3K79me2 show directional preferences in 72% and 71% of the tissues. Additional 12 marks show directional preferences in 50%-70% of the tissues. *These results indicate that active promoters have a directional chromatin signature*.

**Table 1.** For each tissue or cell type, promoters were separated according to the strand to positive and negative groups. A promoter is represented by the 400-nucleotides-long region centered on the transcription start site. Then the region of a promoter was expanded by 2000 nucleotides on each end. Mark vectors over the upstream and the downstream thirds of the expanded regions on the positive strand were compared. A mark is considered directional if these two vectors are tending to be opposite to one another (a negative dotsim value). Not all marks were determined for all tissues. The number of tissues/cell types, for which a mark was determined, is listed under the column titled “Known.” The number of tissues/cell types, in which a mark has directional preference around the promoter regions, is listed under the column titled “Directional.” The percentage of times a mark showed directional preference is listed under the column labeled with “Percentage.” Only marks that were determined for at least five tissues were considered. Very similar results (not shown) were observed on the promoters on the negative strand.

| Mark | Known | Directional | Percentage (%) |
| --- | --- | --- | --- |
| H3K4me3 | 57 | 41 | 72 |
| H3K79me2 | 14 | 10 | 71 |
| H3K4me2 | 16 | 11 | 69 |
| H2AK5ac | 6 | 4 | 67 |
| H3K18ac | 6 | 4 | 67 |
| H2A.Z | 14 | 9 | 64 |
| H3K4me1 | 57 | 35 | 61 |
| H2BK12ac | 5 | 3 | 60 |
| H3K14ac | 5 | 3 | 60 |
| H3K9ac | 24 | 13 | 54 |
| H2BK5ac | 6 | 3 | 50 |
| H3K23ac | 6 | 3 | 50 |
| H3K4ac | 6 | 3 | 50 |
| H3K79me1 | 6 | 3 | 50 |
| H3K27ac | 49 | 22 | 45 |
| H4K91ac | 5 | 2 | 40 |
| H4K8ac | 6 | 2 | 33 |
| H2BK120ac | 6 | 1 | 17 |
| H4K20me1 | 12 | 2 | 17 |
| H3K36me3 | 57 | 6 | 11 |

### Case study: The signatures of high- and low-CpG promoters

It has been reported in the literature that the chromatin signature of high-CpG promoters is different from the signature of low-CpG promoters [47]. In this case study, we use HebbPlot to demonstrate this phenomenon. To this end, we divided promoters active in skeletal muscle myoblasts cells into high-CpG and low-CpG groups using the method proposed by Saxonov, et al. [67]. The high-CpG group consists of 12825 promoters and the low-CpG group consists of 2712 promoters. After that, we generated two HebbPlots from these two groups (Figure 5). These two signatures are very different. Generally, the HebbPlot of the high-CpG group is brighter than that of the low-CpG group, indicating that these histone marks are consistently distributed around the high-CpG promoters. Few marks distinguish the two signatures. The high-CpG group is characterized by the presence of H3K4me3, H3K9ac, and H3K27ac, which are very weak or absent from the low-CpG promoters. The low-CpG group is characterized by the presence of H3K36me3, which is absent from the high-CpG promoters. These two signatures are different from those reported by Karlic, et al. [47]. Two factors may cause these differences. First, the size of the promoter region differs between the two studies. In our study, the size of the promoter is 400 base pairs, while it is defined as 3500 base pairs long (-500 to +3000) in the other study. This longer region is likely to overlap with untranslated and coding regions, whereas it is less likely that the 400-base-pair-long promoter to overlap with these regions. The second factor is that the other study focuses on the correlation between histone marks and expression level, whereas the main purpose of our case study is to visualize the signature of the promoters. Therefore, we believe that our definition is more relevant to the visualization task.

**Figure 5.**
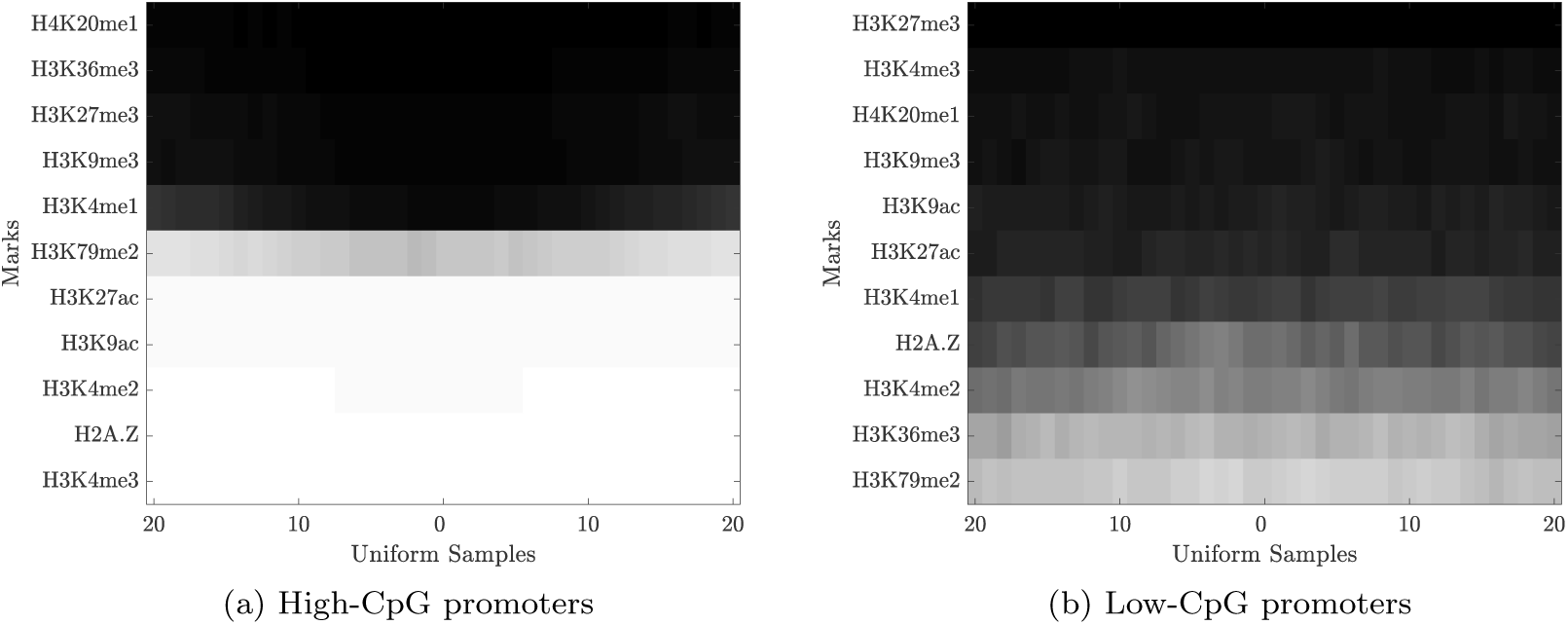
Promoters active in skeletal muscle myoblasts cells were separated into high- and low-CpG groups. A HebbPlot was generated from each group. Clearly, the two signatures are different. Specifically, H3K4me3, H3K9ac, and H3K27ac are present around the high-CpG promoters, whereas they are very weak or absent from the low-CpG promoters. In contrast, H3K36me3 is absent from the high group, but present around the low-CpG promoters. In general, marks present around the high-CpG promoters are stronger than those present around the negative ones.

Next, we performed quantitative comparisons to see if these marks are distributed differently around high- and low-CpG promoters in a consistent way in the 57 tissues. A main advantage of HebbPlots is that they can be compared quantitatively. Comparing two HebbPlots using the dotsim function is simple and easy to interpret; the closer the dotsim value to 1 (-1) the more similar (dissimilar) the two signatures. HebbPlots were generated from the high-CpG promoters (Additional file 4) and the low-CpG promoters (Additional file 5) in the 57 cell types/tissues. We calculated the average dotsim of the two vectors representing a mark around high- and low-CpG promoters in the 57 tissues. Table 2 shows the results. These results confirm that H3K4me3, H3K9ac, and H3K27ac are consistently different around high- and low-CpG promoters (average dotsim value < −0.5). However, H3K36me3 is not different overall (average dotsim value of 0.65). Further, this analysis reveals that H2BK120ac and H4K91ac are also distributed differently around the two groups (average dotsim < −0.5); their signals are stronger around the high-CpG group than the low group.

**Table 2.**
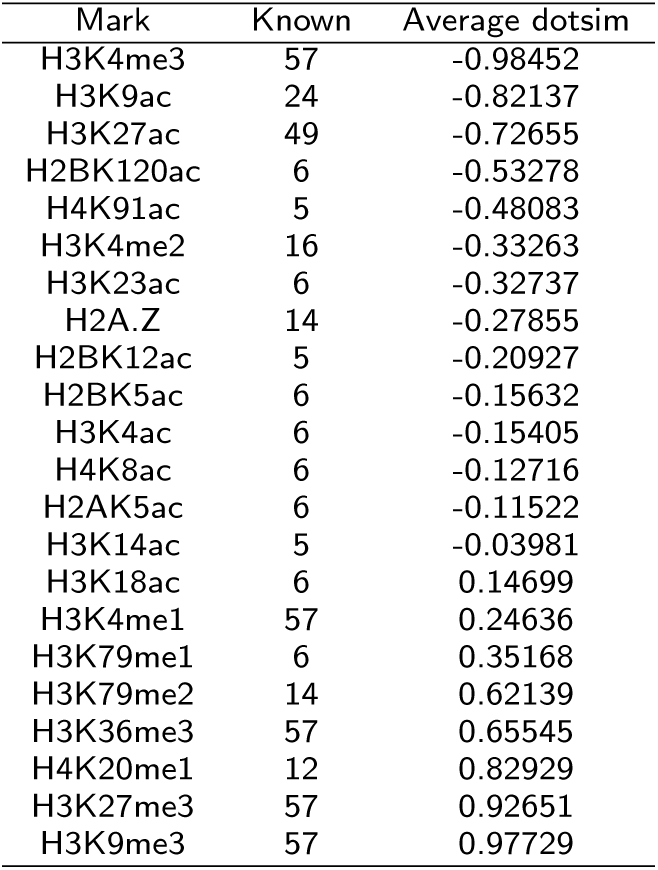
High-CpG promoters have a different signature from that of low-CpG promoters. Active promoters in 57 tissues/cell types were divided into two groups according to their CpG contents. Then two networks were trained on the two groups, producing two signatures for each tissue/cell type. The two signatures of a mark in the same tissue were compared using the dotsim function. The average dotsim values are listed under “Average dotsim.” Not all marks were determined for all tissues. The number of tissues/cell types, for which a mark was determined, is listed under the column titled “Known.”

These results show that the chromatin signatures of high- and low-CpG promoters are different. Specifically, five marks (H3K4me3, H3K9ac, H3K27ac, H2BK120ac, and H4K91ac) are present around high-CpG promoters, whereas they are absent from or very weak around low-CpG promoters.

### Case study: Signature of active enhancers

Here, we demonstrate HebbPlot’s applicability to visualizing the chromatin signatures of enhancers in multiple tissues. To this end, we collected active enhancers from two sources. Enhancers active in H1 (5899 regions) and IMR90 (14073 regions) were obtained from a study by Rajagopal, et al. [54]. Enhancers active in other six tissues were obtained from the Fantom Project. We selected these tissues because they were common to the Fantom and the Roadmap Epigenomics Projects. These enhancers include 5005 regions for liver, 1476 regions for foetal brain, 5991 regions for foetal small intestine, 1619 regions for left ventricle, 11003 regions for lung, and 2225 regions for pancreas.

Next, we generated a HebbPlot from the enhancers of each tissue/cell type. Additional file 6 and Figure 6 show the eight HebbPlots. The HebbPlots of the enhancers active in H1 and IMR90, for which more than 20 marks have been determined, show that multiple marks are abundant around enhancer regions. Similar to what has been reported in the literature, we observed that H3K4me1 is usually stronger than H3K4me3 around enhancers [69]; however there are some exceptions, e.g. foetal brain and lung. H3K27ac and H3K9ac are also present around enhancers, but H3K9me3, H3K27me3, and H3K36me3 are very weak or absent from enhancers. *Further, these HebbPlots suggest that the chromatin signatures of enhancers active in different tissues are similar; however, they are not identical*. For example, H3K27ac is the predominant mark around lung enhancers; H3K4me1 and H3K4me3 are also present, but their signals are weak. In contrast, H3K27ac and H3K4me1 have comparable signals, which are stronger than H3K4me3, around enhancers of foetal small intestine.

**Figure 6.**
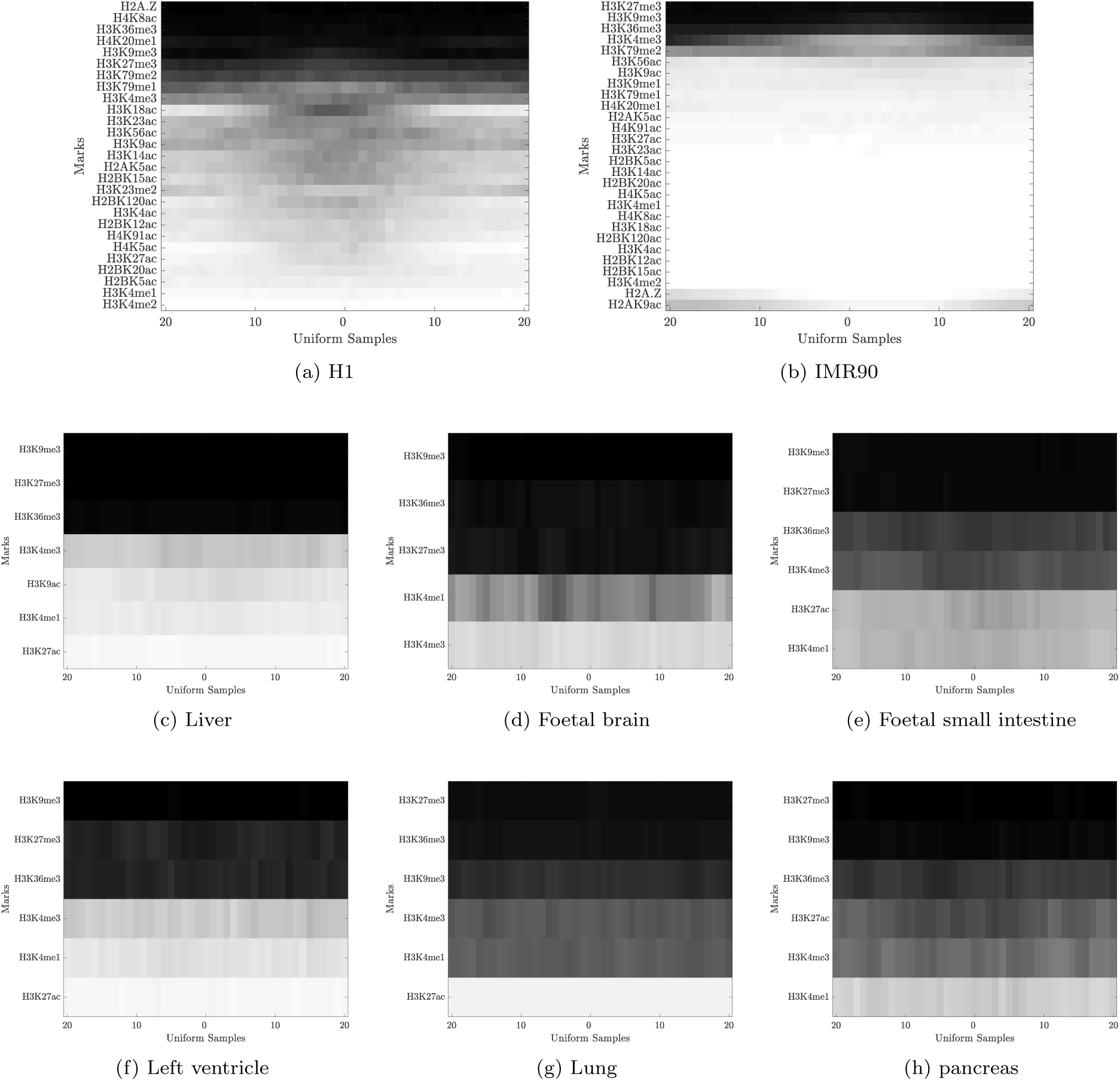
Signatures of active enhancers. Enhancers were collected from a study by Rajagopal et al. [54] and from the Fantom Project. A HebbPlot was generated from the enhancers of each tissue. The HebbPlots of H1 and IMR90, for which more than 20 marks are known, show that several marks are present around active enhancers. Usually, H3K4me1 has a stronger signal around enhancers than H3K4me3; however there are some exceptions, e.g. foetal brain. H3K9ac and H3K27ac are present around enhancers, but H3K9me3, H3K27me3, and H3K36me3 are very weak or absent from enhancers. These plots show that chromatin signatures of enhancers active in different tissues are similar, but not identical.

### Case study: signatures of coding regions of active and inactive genes

Multiple studies indicate that histone marks are associated with gene expression levels [52,70,71]. In this case study, we demonstrate the usefulness of HebbPlot in identifying histone marks associated with high and low expression levels. To start, we divided all genes into nine groups based on their expression levels in IMR90. Then, we generated a HebbPlot from the coding regions of each of these groups (Figure 7). *We found that H3K36me3 and H3K79me1 clearly mark the top two groups. On the lowest six groups, which represent coding regions of inactive genes, these two marks are absent, whereas H3K27me3 is present*. H2A.Z is present in all groups. Generally, the brightness of a HebbPlot decreases as the gene expression levels decrease. In sum, these nine HebbPlots can clearly help with identifying histone marks that are associated with coding regions of active and inactive genes.

**Figure 7.**
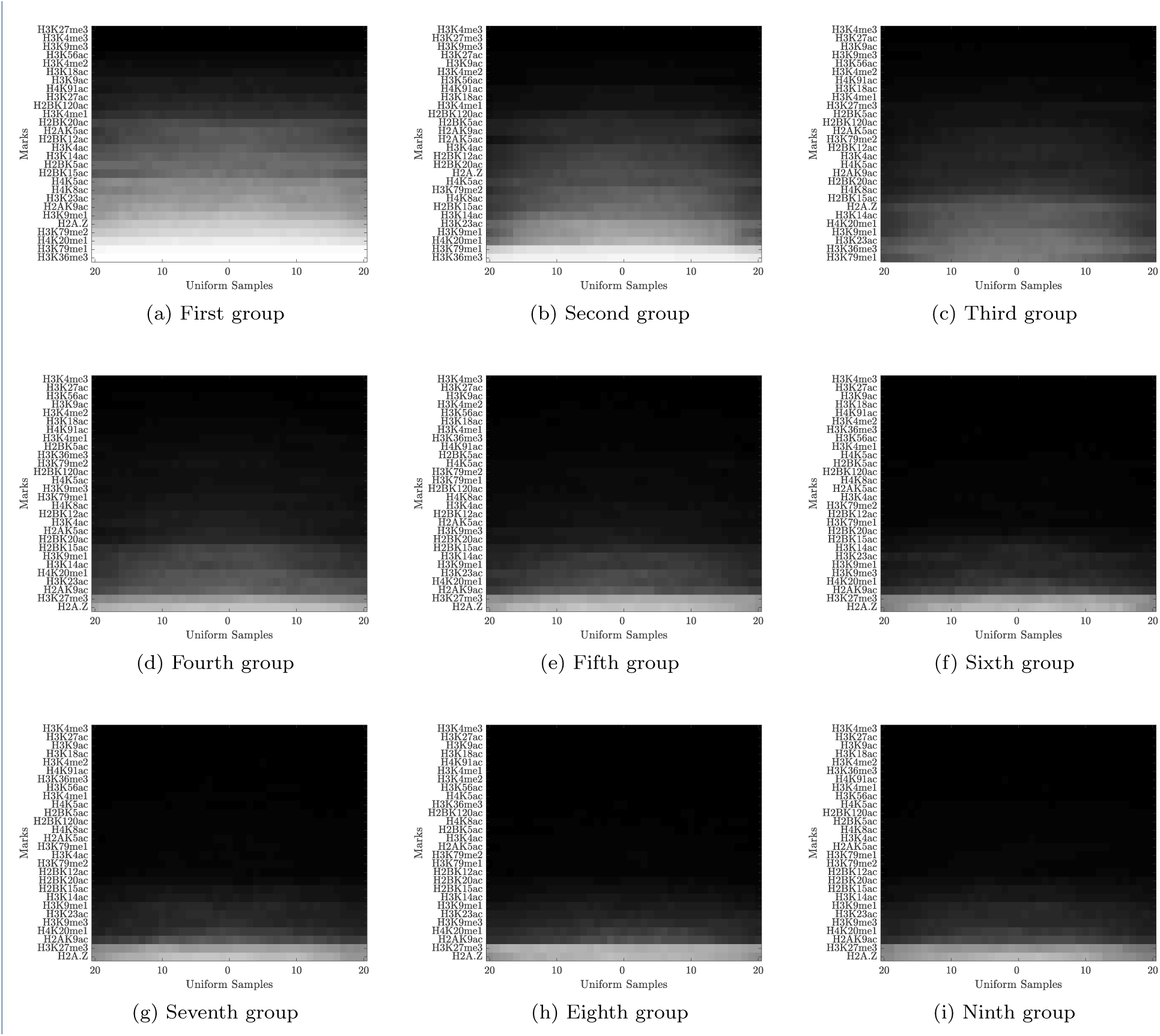
Histone marks are highly associated with gene expression levels in IMR90. Genes were divided into nine groups according to their expression levels. A HebbPlot was generated from the coding regions of each group. In general, the brightness of a HebbPlot decreases as the expression level decreases. The brighter a row, the more consistent its mark is distributed around the set of regions. Further, H3K36me3 and H3K79me1 mark the coding regions of active genes in IMR90 clearly, whereas the repressive modification, H3K27me3, marks the inactive coding regions. H2A.Z is ubiquitous.

After that, we asked whether these marks consistently mark active and inactive coding regions in other tissues/cell types. To answer this question, we generated HebbPlots of coding regions of active (Additional file 7) and inactive (Additional file 8) genes in the 57 tissues/cell types. Then we compared the signatures of each mark around active and inactive coding regions of 57 cell and tissue types using the dotsim function. Recall that dotsim values range between −1 (most dissim ilar) and 1 (most similar). We calculated the average dotsim values of each mark in the 57 tissues/cell types, for which this mark has been determined. Table 3 shows that H3K36me3 and H3K79me1 are very different around active and inactive coding regions (dotsim: −0.86 and −0.64). H3K27me3 is also different (dotsim: 0.44), but the difference is not as strong as H3K36me3 and H3K79me1. After that we asked what other marks are distributed differently around coding regions of active and inactive genes. We found that H3K79me2 consistently marks active coding regions (dotsim: −0.38). Additionally, we found H4K8ac weakly marks active coding regions. Regarding the marks of inactive coding regions, H4K12ac was found to mark these regions (this mark has been determined for one tissue only), whereas H4K14ac and H2AK5ac were found to weakly mark inactive coding regions.

**Table 3.** Histone marks distributions around coding regions of active and inactive genes in 57 tissues. The active and the inactive signatures of a mark in the same cell type are compared using dotsim. The closer the value to −1 (1) is, the more dissimilar (similar) the two signatures are. Because we are interested in dissimilar marks, we only considered marks that have average dotsim of less than 0.5.

| Mark | Average dotsim score | Cell types | Activity |
| --- | --- | --- | --- |
| H3K36me3 | -0.85927 | 57 | Active |
| H4K12ac | -0.67311 | 1 | Inactive |
| H3K79me1 | -0.64466 | 6 | Active |
| H3K79me2 | -0.38126 | 14 | Active |
| H3K14ac | 0.33704 | 5 | Inactive |
| H3K27me3 | 0.43669 | 57 | Inactive |
| H4K8ac | 0.44978 | 6 | Active |
| H2AK5ac | 0.46235 | 6 | Inactive |

*In sum, H3K36me3, H3K79(me1,me2), and H4K8ac mark active coding regions, whereas H4K12ac, H3K14ac, H3K27me3, and H2AK5ac mark inactive coding regions. Generally, the active marks are stronger than the inactive marks*.

### Toward a functional catalog of histone marks

Here, we summarize the findings of our study.

*H2A.Z*

- Directional around promoters stretching upstream.

*H2AK5ac*

- Directional around promoters stretching downstream.
- Weakly associated with coding regions of inactive genes.

*H2BK5ac*

- Directional around promoters stretching downstream.

*H2BK12ac*

- Directional around promoters stretching downstream.

*H2BK120ac*

- Associated with high-CpG promoters.

*H3K4ac*

- Directional around promoters stretching downstream.

*H3K4me1*

- Directional around promoters stretching downstream.
- Absent around transcription start sites.
- Associated with enhancers.

*H3K4me2*

- Directional around promoters stretching downstream.
- Associated with enhancers.

*H3K4me3*

- Directional around promoters stretching downstream.
- Associated with high-CpG promoters.
- Associated with enhancers; however, usually weaker than H3K4me1.

*H3K8ac*

- Weakly associated with coding regions of active genes.

*H3K9ac*

- Directional around promoters stretching downstream.
- Associated with high-CpG promoters.
- Associated with enhancers.

*H3K9me3*

- Weakly associated with coding regions of inactive genes.
- Very weak/absent from enhancers.
- Very weak/absent from promoters.

*H3K14ac*

- Directional around promoters stretching downstream.
- Weakly associated with coding regions of inactive genes.

*H3K18ac*

- Directional around promoters stretching downstream.

*H3K23ac*

- Directional around promoters stretching downstream.

*H3K27ac*

- Associated with high-CpG promoters.
- Associated with enhancers.

*H3K27me3*

- Weakly associated with coding regions of inactive genes.
- Very weak/absent from enhancers.
- Very weak/absent from promoters.

*H3K36me3*

- Associated with coding regions.
- Very weak/absent from enhancers.

*H3K79me1*

- Directional around promoters stretching downstream.
- Associated with coding regions of active genes.

*H3K79me2*

- Directional around promoters stretching downstream.
- Associated with coding regions of active genes.

*H4K12ac*

- Associated with coding regions of inactive genes (this mark is known in one tissue only).

*H4K91ac*

- Associated with high-CpG promoters.

Up to this point, we demonstrated the usefulness of HebbPlot in six case studies. Then we summarized the findings of our study. Next, we discuss the similarities and the differences between HebbPlot and other visualization tools.

## Discussion

Visualization of chromatin marks and their associations with thousands of elements active in a specific cell type is critical to deciphering the function(s) of these marks. Extracting trends and patterns by mere inspection is essentially impossible given that there are more than 100 known chromatin marks and thousands of sequences. As such, it is vital for biologists to have visualization tools to aid in these evaluations. To this end, several tools, Chromatra, ChAsE, and DGW have been developed. In addition, we have created our own visualization technique, HebbPlot. Unlike the other three tools, which cluster genomic regions according to histone modifications, HebbPlot uses an artificial neural network to summarize the data in a form that is convenient for biologists. The following is a brief discussion about HebbPlot and its characteristics that differ from the aforementioned utilities.

Chromatra is a visualization tool that displays chromatin mark enrichment of subregions of each of the input regions. Since it is a plug-in for the well-supported Galaxy platform, it is simple for a user to add it to his or her list of tools. Additionally, this tool is comprised of two modules for chromatin mark analysis. The first module calculates the enrichment scores of a given chromatin mark to a given set of genomic locations of interest. The second module, while similar to the first, adds the additional functionality of clustering the results by additional parameters, e.g. gene expression levels. All results of these modules are then projected onto a heat map, which can be exported for further research. While Chromatra’s ease-of-use and versatility are common characteristics between it and HebbPlot, HebbPlot takes a dramatically different approach to how it clusters data. Whereas Chromatra handles enrichment levels in genomic regions of variable length through binning, HebbPlot will extract the same number of points for any region. HebbPlot will then utilize an artificial neural network to derive a representative pattern for the chromatin marks across all of the points in every region. Our tool proceeds to cluster the patterns for each chromatin mark according to their similarity to each other, and then produces a heatmap of the results. Therefore, rather than evaluate genomic regions that have been mapped to chromatin marks, HebbPlot summarizes the distribution of each chromatin mark across a “representative” region. This allows researchers to only have to view one heatmap before acquiring a solid understanding of how the histone modifications are represented across the regions.

ChAsE and HebbPlot have their basis in displaying information clearly and easily to the user. Their design philosophy is rooted in the fact that many visualization tools demand a high amount of technical knowledge that is unreasonable to expect from researchers. With this said, HebbPlot and ChAsE also diverge significantly in how they cluster the input and how they present their results. Similar to Chromatra, ChAsE will cluster regions together based on the abundance of chromatin marks (or any genomic area of interest) in each region. Afterwards, ChAsE allows the user the flexibility to inspect the clusters further via methods like K-Means clustering and signal queries. HebbPlot, as explained before, samples a fixed number of points in each given region of interest. These samples, and the overlapping marks, are then processed by an artificial neural network to determine a motif for each histone modification that is illustrative of its distribution in all given regions. The motifs for each considered modification is then clustered in a hierarchy so that all modifications of similar enrichment levels are placed together. A figure of this detailed clustering is then produced, providing researchers with a way to quickly understand how histone mark abundance is distributed across the locations.

DGW is a tool that consists of two modules. The first is an alignment and clustering module, whereas the second is a visualizer for the results. DGW is designed to “rescale and align” the histone marks of genomic regions (such as TSSs and splicing sites). Additionally, it hierarchically clusters the aligned marks into distinct groups. Regarding the visualization module, DGW creates heat maps and dendrograms of chromatin marks of a set of genomic locations. Similar to our conclusions in the previous comparisons, there are several notable similarities and differences between DGW and HebbPlot. HebbPlot is similar to DGW in that it scales the regions. However, HebbPlot implements it using a different idea. Specifically, HebbPlot samples a fixed number of equally spaced points from each region regardless of the region length. HebbPlot learns a general pattern of chromatin marks summarizing all of the input regions as one representative region. Unlike DGW, hebbPlot does not cluster the input regions based on the distribution of a mark. Hierarchical clustering is utilized in HebbPlot not to cluster the regions according to the enrichment of a mark, but to cluster all marks according to their distributions around the representative region. The amount of details produced by DGW can be inappropriate in the presence a large number of marks and regions. HebbPlot on the other hand, is built specifically to make large amounts of data manageable and meaningful for biologists through its summarization technique.

Our comparisons regarding these four tools makes it clear that the advantages provided by HebbPlot are not well represented among related tools. There are numerous tools for clustering regions according to the abundance of chromatin marks, but besides conventional plots, there are hardly any techniques for determining the patterns of marks across all regions. This means it is important for HebbPlot to coexist among other popular visualization tools. Its unique and concise summarization of data is vital to evaluating a large number of chromatin signals across thousands of regions. This is not to say that the level of description provided by other tools is not useful. Indeed, biologists need to be able to see the specific results that other utilities facilitate. However, what HebbPlot offers is a look at the “big picture” of the data, giving the biologist an easy way to understand how each chromatin mark is distributed across a large number of regions.

## Conclusions

In this manuscript, we described a new software tool, HebbPlot, for learning and visualizing the chromatin signature of a genetic element. HebbPlot produces a simple image that can be interpreted easily. Further, signatures learned by HebbPlot can be compared quantitatively. We validated HebbPlot in six case studies using 57 human tissues and cell types. The results of these case studies are novel or confirming to previously reported results in the literature, indicating the accuracy of HebbPlot. We found that active promoters have a directional chromatin signature; specifically, H3K4me3 and H3K79me2 tend to stretch downstream, whereas H2A.Z tends to stretch upstream. Further, our results confirm that high-CpG and low-CpG promoters have different chromatin signatures. When we compared the signatures of enhancers active in eight tissues/cell types, we found that they are similar, but not identical. Contrasting the signatures of coding regions of active and inactive genes reveals that certain modifications (H3K36me3, H3K79me1, H3K79me2, and H4K8ac) mark active coding regions, whereas different modifications (H4K12ac, H3K14ac, H3K27me3 and H2AK5ac) mark coding regions of inactive genes. Our study results in a visual catalog of chromatin signatures of multiple genetic elements in 57 human tissues and cell types. Further, we made a progress toward a functional catalog of more than 20 histone modifications. Finally, HebbPlot is a general tool that can be applied to a large number of studies, facilitating the deciphering of the histone code.

## Availability and requirements

The source code (Perl and Matlab) is available as Additional file 1.

**Project name:** HebbPlot.

**Project home page:** https://github.com/TulsaBioinformaticsToolsmith/HebbPlot

**Operating system(s):** UNIX/Linux/Mac.

**Programming language:** Perl and Matlab.

**Other requirements:** Matlab Statistics and Machine Learning Toolbox and Bedtools (http://bedtools.readthedocs.io/en/latest/).

**License:** Creative Commons license (attribution + non-commercial + no derivative works).

**Any restrictions to use by non-academics:** License needed.

## Author’s contributions

HZG designed the software and the case studies, implemented the neural network, and wrote the manuscript. AV coded the software, processed the data, and wrote the manuscript. ZER processed the data and wrote the manuscript.

## Acknowledgements

The authors would like to thank Michael Buck, Associate Professor of biochemistry at the State University of New Your at Buffalo, for useful discussions. This research was supported by internal funds provided by the College of Engineering and Natural Sciences and the Faculty Research Grant Program at the University of Tulsa.

## Additional Files

Additional file 1 — HebbPlot Software

This compressed file (.tar.gz) includes the source code (Matlab and Perl) of HebbPlot.

Additional file 2 — HebbPlots of active promoters on the positive strand

This compressed file (.tar.gz) include HebbPlots of promoters on the positive strand active in 57 tissues/cell types.

Additional file 3 — HebbPlots of active promoters on the negative strand

This compressed file (.tar.gz) include HebbPlots of promoters on the negative strand active in 57 tissues/cell types.

Additional file 4 — HebbPlots of high-CpG promoters

This compressed file (.tar.gz) include HebbPlots of high-CpG promoters active in 57 tissues/cell types.

Additional file 5 — HebbPlots of low-CpG promoters

This compressed file (.tar.gz) include HebbPlots of low-CpG promoters active in 57 tissues/cell types.

Additional file 6 — HebbPlots of active enhancers

This compressed file (.tar.gz) include HebbPlots of enhancers active in eight tissues/cell types.

Additional file 7 — HebbPlots of coding regions of active genes

This compressed file (.tar.gz) include HebbPlots of genes active in 57 tissues/cell types.

Additional file 8 — HebbPlots of coding regions of inactive genes

This compressed file (.tar.gz) include HebbPlots of genes inactive in 57 tissues/cell types.

